# Pan-cancer analysis on microRNA-mediated gene activation

**DOI:** 10.1101/462960

**Authors:** Hua Tan, Shan Huang, Zhigang Zhang, Xiaohua Qian, Peiqing Sun, Xiaobo Zhou

**Affiliations:** School of Biomedical Informatics, The University of Texas Health Science Center at Houston, Houston, TX 77030, USA; Department of Cancer Biology, Wake Forest Comprehensive Cancer Center, Wake Forest School of Medicine, Winston-Salem, NC 27157, USA; Institute for Medical Imaging Technology, School of Biomedical Engineering, Shanghai Jiao Tong University, Shanghai 200030, China

**Keywords:** pan-cancer miRNA, miRNA activation, intragenic miRNA, super-enhancer

## Abstract

While microRNAs (miRNAs) were widely considered to repress target genes at mRNA and/or protein levels, emerging evidence from *in vitro* experiments has shown that miRNAs can also activate gene expression in particular contexts. However, this counterintuitive observation has rarely been reported or interpreted in *in vivo* conditions. We systematically explored the positive correlation between miRNA and gene expressions and its potential implications in tumorigenesis, based on 8375 patient samples across 31 major human cancers from The Cancer Genome Atlas (TCGA). Results indicated that positive miRNA-gene correlations are surprisingly prevalent and consistent across cancer types, and show distinct patterns than negative correlations. The top-ranked positive correlations are significantly involved in the immune cell differentiation and cell membrane signaling related processes, and display strong power in stratifying patients in terms of survival rate, demonstrating their promising clinical relevance. Although intragenic miRNAs generally tend to co-express with their host genes, a substantial portion of miRNAs shows no obvious correlation with their host gene due to non-conservation. A miRNA can upregulate a gene by inhibiting its upstream suppressor, or shares transcription factors with that gene, both leading to positive correlation. The miRNA/gene sites associated with the top-ranked positive correlations are more likely to form super-enhancers compared to randomly chosen pairs, suggesting a potential epigenetics mechanism underlying the upregulation. Wet-lab experiments revealed that positive correlations partially remain in the *in vitro* condition. Our study provides the field with new perspectives on the critical role of miRNA in gene regulation and novel insights regarding the complex mechanisms underlying miRNA functions, and reveals the clinical significance of the potential positive regulation of gene expression by miRNA.

## INTRODUCTION

Cancer is caused by uncontrolled cell growth reflective of multiple established hallmarks (Hanahan and Weinberg, 2011). Underlying the aberrant cell proliferation is activation of critical oncogenes and inactivation of tumor suppressor genes resulted from multiple genetic and epigenetic alterations in a cancer tissue-specific manner (Dawson and Kouzarides, 2012;Tan et al., 2012, 2015;Vogelstein et al., 2013). Among these, microRNAs (miRNAs) are a class of small (~22 nucleotides), non-protein-coding RNAs known as important post-transcriptional regulators of gene expression (Bartel, 2004;He and Hannon, 2004). MicroRNAs exert regulatory functions by base-pairing with complementary sequences typically in the 3’-untranslated region (3’UTR) of mRNAs to target them for degradation or prevent their translation (Bartel, 2004, 2009). It is estimated that more than 60% of human protein-coding genes are under selective pressure to maintain pairing to miRNAs and over one third of human genes appear to be conserved miRNA targets (Friedman et al., 2009;Lewis et al., 2005). This indicates that miRNAs can influence almost every critical signaling pathway in a cell (Filipowicz et al., 2008).

A canonical miRNA is transcribed from miRNA gene by RNA polymerase II (pol II) as a primary miRNA (pri-miRNA). The pri-miRNA transcript is first cleaved in the nucleus by the microprocessor, which contains a nuclear RNase III called Drosha and its cofactor DGCR8, into a hairpin structured precursor miRNA (pre-miRNA). Then, the pre-miRNA is exported to the cytoplasm through the activation of Exportin 5 and RAN-GTP, and further processed by an endonuclease Dicer to generate the miRNA duplex, which contains the miRNA paired to its passenger strand usually called miRNA*. One strand of the miRNA duplex, the mature miRNA, is loaded into an Argonaute protein (AGO) to form a miRNA-induced silencing complex (miRISC), whereas the other strand, the miRNA*, is degraded. Once loaded into the silencing complex, the miRNA pairs to complementary sites within mRNAs or other transcripts and the Argonaute protein exerts the posttranscriptional repression (Bartel, 2018). Considering that ~2000 miRNA gene have been identified in human genome (Kozomara and Griffiths-Jones, 2014), that a miRNA can target hundreds even thousands of different mRNAs, and an individual mRNA might be influenced by multiple miRNAs (Agarwal et al., 2015), the miRNA biogenesis pathway plays an essential role in shaping the gene regulatory networks. Therefore, the miRNA biogenesis is elaborately maintained in a favorable balance under physiological condition, but can be severely impaired in cancer, resulting in differential expression of critical miRNAs compared to that in the normal tissues (Croce, 2009;Volinia et al., 2006).

Based on the canonical miRNA biogenesis pathway and mechanism of function, miRNAs have long been believed to elicit their effects only through mRNA degradation and/or translation inhibition. Recently, evidence has accumulated to suggest that miRNA can also promote gene expression in particular contexts via various mechanisms. For instance, under cell cycle arrest, human miRNA miR-369-3 was reported to activate tumor necrosis factor-α (TNFα) translation by directing micro-ribonucleoproteins (microRNPs) including AGO and FXR1 to a special type of miRNA binding sites called AU-rich elements (AREs) in the 3’UTR of TNFα, while it was also acknowledged that this translational activation can be switched to repression in proliferating cells (Vasudevan et al., 2007). In most cases, however, miRNA was reported to activate gene expression by binding to the promoter region followed by recruitment of transcription factors to the miRNA binding sites. Striking examples include: miR-373 activates E-cadherin (CDH1) and cold-shock domain-containing protein C2 (CSDC2) by binding to their promoter regions (Place et al., 2008), miR-205 induces the expression of tumor suppressor genes interleukin 24 (IL24) and IL32 by targeting specific sites in their promoters (Majid et al., 2010), miR-744/1186/466d-3p induces Ccnb1 expression in mouse cell lines by targeting promoter elements (Huang et al., 2012), miR-589 binds the promoter RNA and activates cyclooxygenase-2 (COX-2) transcription (Matsui et al., 2013), and let-7i binds to TATA-box of IL2 and activates it at both mRNA and protein level (Zhang et al., 2014). Recently, a new mechanism regarding miRNA activation has been proposed, showing that miR-24-1 activates FBP1 and FANCC genes by targeting their enhancers (Xiao et al., 2017). In addition, these studies invariably corroborated that proteins related to miRNA biogenesis or functions, such as AGO and Dicer, or transcription related enzymes are significantly enriched surrounding the binding sites during gene activation. Collectively, these previous findings, mainly observed in *in vitro* conditions, established that miRNA can also upregulate gene expression by directly binding to the transcriptional regulatory regions.

While the miRNA-mediated gene activation has been well studied in *in vitro* conditions and in animal experiments, it was rarely explored in human samples. To address the gap, we leveraged the Cancer Genome Atlas (TCGA) and conducted an unprecedentedly comprehensive analysis on the miRNA-gene interaction profiles in 8375 patient samples across 31 major human cancer types. We checked the correlations between all 1046 miRNAs and 20531 genes annotated in TCGA, and found that positive miRNA-gene correlation is surprisingly prevalent and consistent across human cancers, even when the gene bears conserved binding sites for the miRNA. And the positive correlations display disparate patterns compared to the negative correlations. We performed a series of stringent bioinformatics analysis to investigate whether this positive correlation has any biological or clinical implication especially in the context of human oncogenesis. We revealed that miRNA and gene pairs with positive correlation are extensively involved in many biological processes pertaining to immune response, cell membrane signaling, cell cycle control and other cancer hallmarks. In addition, the top ranked miRNAs and genes can well stratify patients in terms of overall survival rate based on their single or combined expression level. These results warrant the biological and clinical significance of the widespread positive correlations between miRNAs and genes across human cancers.

We further investigated the molecular mechanisms underlying the observed miRNA-gene positive correlation. Most of the positive correlations (~87%) can be explained by one or more of our proposed four indirect-regulation hypotheses, including miRNA-host gene co-expression, gene activation by inhibiting upstream suppressor, co-regulation by shared transcription factors, and co-activation by common histone modifications. Considering that co-expression by shared genetic or epigenetic factors cannot be viewed as causative relationship, our study stresses that although some positive correlations are implicated in tumorigenesis, the expression level of the miRNA and gene are independent on each other. On the other hand, mechanisms related to the miRNA-host gene co-expression and indirect activation of gene by inhibiting its upstream suppressor involve causation, in the sense that the expression level of one will influence that of the other. We further hypothesized that the remaining part of positive correlations (~13%) can be explained by direct binding of miRNA to particular transcriptional regulatory regions of the partner gene, as reviewed above. Our wet-lab experiments in corresponding cancer cell line only partially recapitulated the positive correlations observed in human patient samples, implying that the miRNA-gene interaction are dramatically different between *in vitro* and *in vivo* conditions. This in turn highlights the significance of a comprehensive analysis on the miRNA-directed gene activation in human samples.

## RESULTS

### Positive miRNA-gene correlations are prevalent and consistent across human cancers

By an integrative miRNA-gene correlation analysis on 1046 miRNAs and 20531 genes across 31 TCGA cancers (Table S1), we detected a total of 2,842,030 pairs that were significantly positively correlated (*R* > 0.1, adj.P < 0.05) in at least one TCGA cancer type. We then ranked these positive pairs according to the number of cancer types (called cancer coverage) in which their correlation appear to be positive (*R* > 0.1) and significant (adj.P < 0.05). In most of the subsequent analysis, we focus on the top ranked pairs with cancer coverage ≥ 10, totaling 18996 miRNA-gene pairs, which involves 348 miRNAs and 3074 genes (Table S2). Each of the top 56 significant positive pairs covered at least 27 TCGA cancer types (Figure 1A). Three miRNA~gene pairs, including miR-196b~HOXA10, miR-335~MEST and miR-483~IGF2, were significantly positively correlated across all 31 TCGA cancers under investigation. Interestingly, miR-196b and HOXA10 have been reported to co-express and their overexpression characterized poor prognosis in patients with gastric cancer (Lim et al., 2013). The IGF2 intronic miR-483 has been widely recognized as an oncogenic miRNA that transcriptionally upregulates its host gene IGF2 (Liu et al., 2013;Veronese et al., 2010), while the long non-coding RNA (lncRNA) H19 intragenic miR-675 was shown to be the most highly conserved feature of H19 and serves as the functional regulatory unit of this lncRNA (Keniry et al., 2012). These previous studies corroborated the significant biological implications of the positive correlations that are prevalent across multiple cancer types.

**Figure 1.**
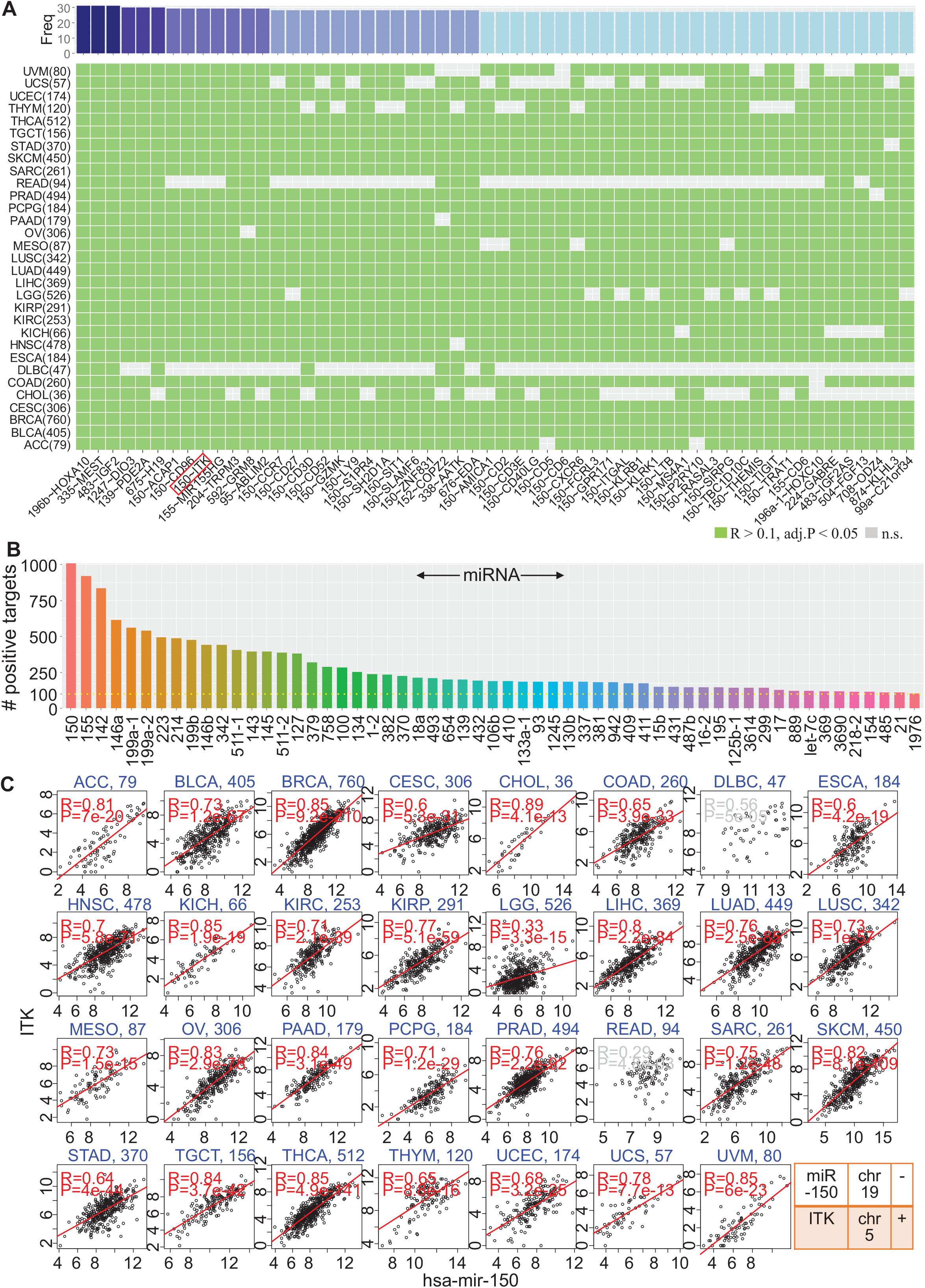
Overview of miRNA-gene positive correlation landscape in human cancers. (A) Heat map showing top positively correlated miRNA-gene pairs covering ≥27 cancer types. Cancer coverage (Freq) for each pair is shown on the top. TCGA cancer names and corresponding sample sizes used for correlation calculation is shown on the left. Pairs are shown in the format of miRNA~gene at the bottom, with human miRNA prefix (hsa-mir-) omitted for better visualization. A positive correlation with Pearson correlation coefficient *R* > 0.1 and Hochberg adjusted p-value < 0.05 was deemed significant. n.s.: non-significant. (B) Top miRNAs ranked by the number of its positively correlated genes. For better visualization, only miRNAs with ≥100 targets and across ≥10 cancer types are shown. (C) Detailed correlation profiles of a representative positive pair hsa-mir-15 ~ITK across 31 TCGA cancers. The sample size of each cancer, Pearson correlation coefficient and p-value are also shown. Red color indicates significant positive correlation. See also Figure S1-S3.

We further extracted the miRNAs and genes from the top-ranked pairs with cancer coverage ≥ 10, mapped each miRNA to all genes positively correlated with it (positive targets), and ranked the miRNAs according to the number of their positive targets. The top 57 miRNAs were found to positively correlate with at least 100 genes (Figure 1B). We also explored the genomic spatial distribution of the positive targets of each miRNA, and found that their positive targets tend to distribute along the whole genome evenly (Figure S2). The top ranked miRNAs can be clustered into several groups based on their positive targets, and miRNAs in the same group tend to share a large number of positively correlated genes (Figure S3). miR-150, a miRNA located at chromosome 19, turns out to be the most active miRNA, which positively correlates with 1009 different genes each covering at least 10 cancers. It is an important hematopoietic cell (especially B-cell, T-cell and natural killer cell)-specific miRNA and has been widely reported to be differentially expressed in cancers especially in hematopoietic malignancies compared to normal controls (He et al., 2014). A detailed correlation profile for a representative pair miR-150 ~ ITK (top 9) shows that they are positively correlated in 29 out of 31 TCGA cancers. Even in the exceptional scenarios in DLBC and READ cancers, miR-150 shows obvious positive correlation with ITK, although was not detected as significant due to the relatively small sample size of these two cancer types and the stringent adjusted p-value threshold in this study. Collectively, these findings demonstrate that significant positive correlation between miRNAs and genes is highly prevalent and consistent across different cancer types.

### Positive miRNA-gene correlations display distinct patterns than negative correlations

Our study revealed several unprecedented patterns of miRNA-gene interactions in human patient samples. First, in comparison to negative correlations, the miRNA-gene positive correlations are much more significant, prevalent and consistent across cancer types (Figure 1 and Figure S1). Using the same thresholds of correlation coefficient and p-value, we detected 18996 miRNA-gene pairs that are positively correlated in at least ten cancer types (PanCan10), compared to 1514 pairs with negative correlation (Figure S1B). And the negative correlations generally cover much less cancer types than the positive ones (Figure S1A, S1D). Interestingly, the constituent miRNAs and genes of these positive and negative pairs are largely overlapped (Figure S1B), suggesting that there might be a constant and small group of miRNAs and genes taking effect in a biological system. We also noted that the positive correlation of about 17% (3223/18996) pairs retained in the normal samples (Figure S1E), indicating that miRNAs can function by upregulating particular genes under physiological besides oncogenic conditions. Second, a miRNA can significantly correlate with multiple genes and vice versa, potentially forming a multiple-to-multiple regulation network, in both positive and negative correlation scenarios. While a miRNA can significantly correlate with an incredibly large number of genes, for example, miR-150 positively correlate with 1009 genes in PanCan10 group and miR-141 negatively correlate with 1140 genes in PanCan7 negative group (Figure 1B and Figure S1C), their correlated partners tend to distribute nearly evenly across the whole genome, instead of clustering in particular genomic regions (Figure S2). Third, miRNAs in the top-ranked positive correlations can be well clustered according to their correlated partner genes. Particularly, the top three miRNAs miR-150/155/142 share 750 partner genes, an overwhelmingly major part of each individual’s partners. Similar pattern exists in the miR-214/199b/199a-2/199a-1 and miR-337/409/431/654 clusters (Figure S3). Together, these results revealed many previously unrecognized miRNA-gene interaction patterns in human cancer, especially the difference between miRNA-gene positive and negative correlations.

### Positive miRNA-gene correlations demonstrate biological and clinical significance

We defined the cancer coverage of a significant miRNA-gene pair as the number of cancer types in which this pair shows positive correlation. To better focus on the top-ranked significant pairs (cancer coverage ≥ 10, Table S2), we categorized the pairs into four groups with increasing pan-levels: PanCan10 (covering ≥10 TCGA cancer types, n = 18996), PanCan15 (n = 4683), PanCan20 (n = 1316) and PanCan25 (n = 240) (Figure 2A, top). To explore the biological and clinical significance of the miRNAs and genes involved in the top-ranked positive pairs, we investigated their implications in cancer initiation and progression from three perspectives. First, we examined the associations of the miRNAs with well-known cancer hallmarks, and the enrichment of the genes in established hallmark gene sets of different important biological processes. Second, we explored the major biological processes that the genes participate by gene ontology (GO) term and KEGG signaling pathway enrichment analysis. Third, we assessed the stratification power of these miRNAs and genes in characterizing the patient overall survival rate, by dividing patients into different groups based on the constituent miRNAs and/or genes of each positive pair (Figure 2A, bottom).

**Figure 2.**
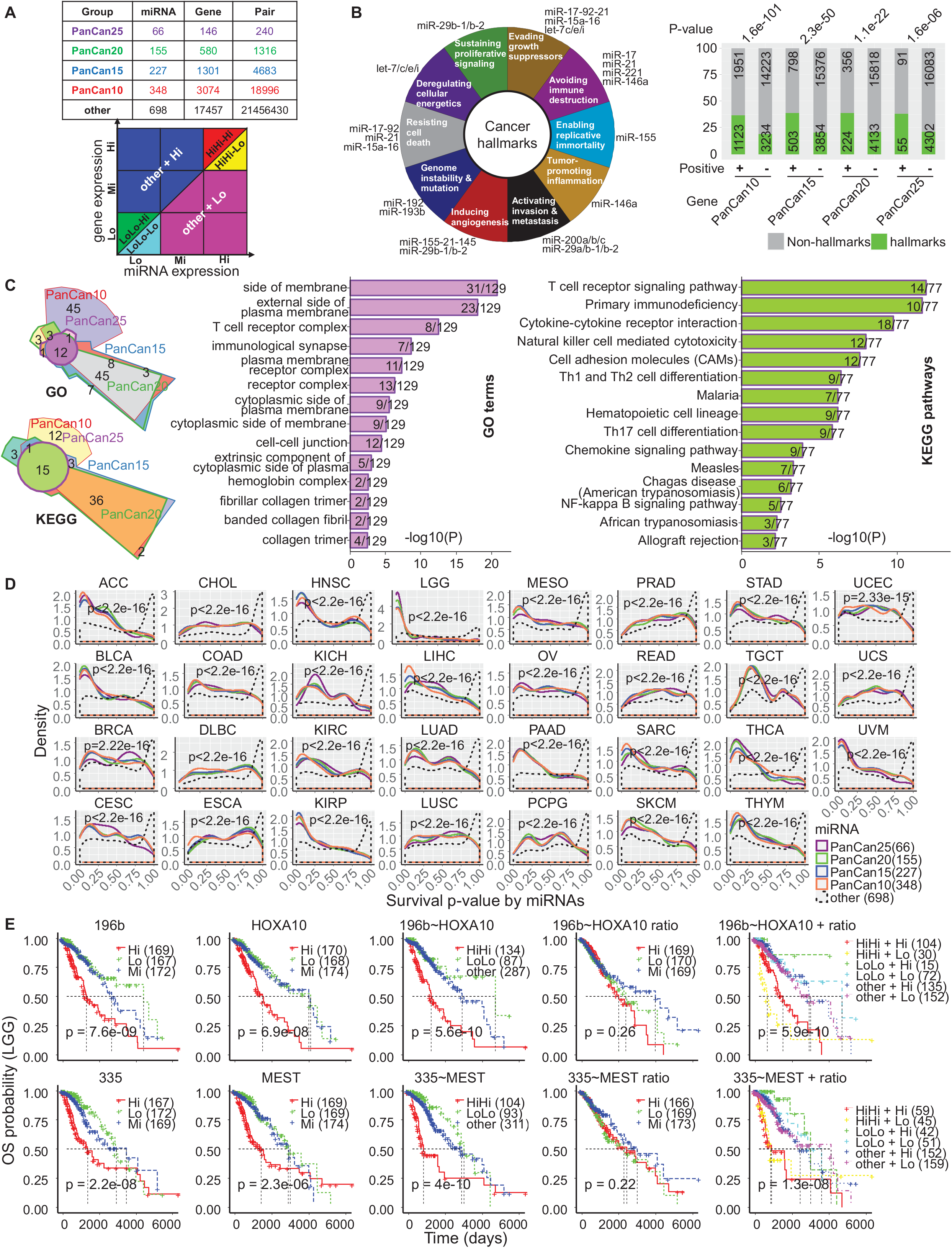
Biological and clinical significance of miRNA-gene positive correlations. (A) Top: Number of miRNAs, genes, and pairs in each pan-level group. The group “other” refers to pairs not covered by PanCan10. Bottom: Patient samples were divided into different groups based on their miRNA and/or gene expression. (B) Left: Association between top-ranked (PanCan10) miRNAs and cancer hallmarks adapted from Hanahna and Weinberg 2011. Right: Enrichment of top-ranked genes of different pan-levels in 50 hallmark gene sets (pooled together) summarized in Liberzon et al. 2015. Numbers inside bars represent the number of genes detected in that group. +/-refers to genes from/not from the top-ranked positive pairs of corresponding pan-levels. P-values were assessed by hypergeometric test. (C) Left: Chow-Ruskey diagrams showing numbers of significant GO terms and KEGG signaling pathways (P<0.05) participated by genes from four pan-level groups as shown in (A). Middle: GO terms shared by genes from the four pan-level groups. Right: KEGG processes shared by genes from the four pan-level groups. (D) Density distributions of p-values of survival analysis (log-rank test) based on expression of miRNAs from indicated pan-level groups. The stratification power in patient survival of miRNAs from different pan-level groups was compared by K-S test. Cancer type and p-value in K-S test are shown on each subgraph. (E) Representative survival analysis results for the top two (ranked by cancer-coverage as in Figure 1A) positive pairs – miR-196b~HOXA10 and miR-335~MEST in lower grade glioma (LGG) cancers. From left to right panels, patients were stratified into different groups as shown in (A) by miRNA only, gene only, miRNA/gene combination, miRNA/gene ratio (gene:miRNA), miRNA/gene combination and ration, followed by a log-rank test in each stratification scheme. See also Figure S4-S5.

We found that all previously reported cancer hallmark miRNAs (Suzuki et al., 2017) were covered by the PanCan10 miRNAs (Figure 2B, left). In addition, genes from all pan-level groups were significantly enriched in the pooled hallmark gene sets of essential biological processes (Liberzon et al., 2015) including cell development, metabolism, immune and proliferation, which are assumedly critical for cancer development and progression (Figure 2B, right; Table S3).

The enriched gene ontology terms and signaling pathways for genes at different pan-levels converged to 14 significant GO terms and 15 significant KEGG pathways (Figure 2C, left; Figure S4). The enriched GO terms primarily consisted of cytoplasmic membrane and various receptor complexes on the membrane (Figure 2C, middle), while the enriched KEGG pathways largely covered the immune signaling related processes, including T cell receptor signaling and T helper cell (Th1, Th2, and Th17) differentiation, natural killer cell cytotoxicity and NF-kappa B signaling pathway (Figure 2C, right).

We used miRNAs from different pan-level groups to stratify patients in terms of overall survival rate, depicted by survival log-rank test p-values. P-value density showed a significant left-shift of p-value in all pan-level miRNA groups across all cancers, compared to the control group – the remaining 698 miRNAs not in any pan-level group (Figure 2D). This indicates that miRNAs in the top-ranked positive pairs represent much better discriminator of patient survival compared to the other miRNAs. This trend is also applicable to the genes from different pan-level groups (Figure S5A). In addition, we noticed that in most cancers combined miRNA and gene expression could significantly improve the stratification power of miRNA or gene alone, while further integration of the gene:miRNA ratio showed stratification performance comparable to the optimal method of all other scenarios (Figure S5B). Take the top two positive pairs miR-196b~HOXA10 and miR-335~MEST in lower grade glioma (LGG) cancer as an example (Figure 2E), the associated miRNA and gene alone can well stratify the patients individually, while combination of the pair further improved the stratification. Specifically, patients with both higher miRNA and gene expression exemplified much poorer prognosis than those with both lower miRNA and gene expression, or all others absent from these two groups, and this distinction is more significant in comparison with the scenarios of miRNA or gene alone. When the gene:miRNA ratio was introduced and patients were further divided into smaller subgroups, the stratification significance remained comparable.

### Intragenic miRNAs tend to but do not always co-express with their host genes

Some miRNAs are intragenic and hence assumedly to be transcribed together with their host transcription unit which can be a protein-coding gene or a long non-coding RNA. This hypothesis has been leveraged to facilitate identification of miRNA target genes, based on the correlation between its candidate targets and host gene (Gennarino et al., 2009). We reasoned that a part of the top-ranked positive correlations detected in our study might be ascribed to this intragenic co-expression hypothesis. We checked the genomic locations of all miRNAs and genes from annotation databases, and obtained 591 miRNAs that are embedded in another gene (host gene), 451 of which are covered in the TCGA data. Hypergeometric test results demonstrated a consistently strong tendency of co-expression between intragenic miRNA and its host gene at all pan-levels (Figure 3A, Table S4). When all pairs covering at least one cancer type were pooled together, the tendency remained significant (P=8×10^-108^). In addition, conserved miRNAs (conservation adopted from TargetScanHuman, see Methods and Table S4) are more likely to co-express with their host genes compared to their non-conserved counterparts (P=5.3×10^-25^), consistent well with previous research (Franca et al., 2016;He et al., 2012).

**Figure 3.**
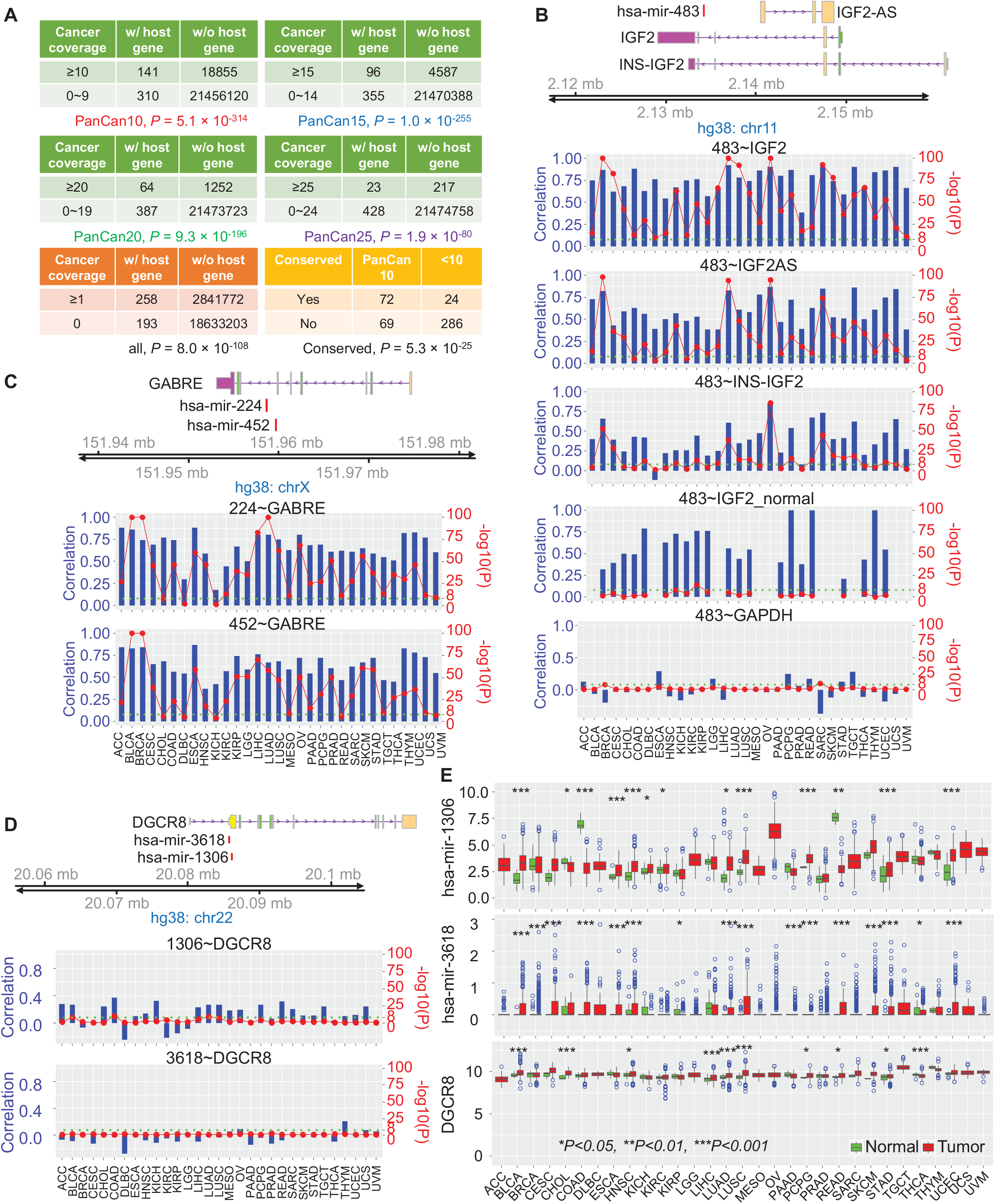
miRNAs tend to co-express with their host genes. (A) Overview of intragenic miRNA-host gene co-expression patterns. The first five tables show the tendency of a miRNA-host gene pair to be positively correlated at each pan-level respectively. Cancer coverage refers to the number of cancer types in which a specific pair is positively correlated. The last table shows the tendency of an intragenic miRNA co-expressing with its host gene to be conserved across species. Conservation was predicted by TargetScanHuman. P-values were assessed by hypergeometric test. (B) Correlation between hsa-mir-483 and its host gene IGF2, isoform of the host gene, INS-IGF2, and IGF2 antisense, IGF2-AS. The correlation between hsa-mir-483 and IGF2 in normal tissues, as well as correlation between hsa-mir-483 and housekeeping gene GAPDH are also shown for comparison and as control. Pearson correlation coefficient and significance level (-log10 transformed) are illustrated by blue bars and red solid circles respectively. A threshold - log10(P)=8 is marked by a green dotted line. Normal tissues which are not available in TCGA are left blank. (C) Correlation between gene GABRE and its two embedding miRNAs: has-mir-224 and hsa-mir-452. Shape and color code are same as (A). (D) Left: Correlation between microprocessor complex subunit DGCR8 and its two embedding miRNAs: hsa-mir-3618 and hsa-mir-1306. Shape and color code are same as (A). (E) Expression level of the two intragenic miRNAs has-mir-1306/3618 and their host gene DGCR8 in 31 TCGA cancer tissues and in comparison with their normal controls if available. The middle line in the box is the median, the bottom and top of the box are the first and third quartiles, the whiskers extend to 1.5 IQR (interquartile range) of the lower quartile and the upper quartile respectively, and the blue hollow circles represent outliers. Expression difference significance (p-value) was assessed by two-sided student *t*-test. See also Figure S6 and Table S4.

As an example, the insulin-like growth factor-2 (IGF2) and its intragenic miRNA, miR-483, which is located on intron 2 of IGF2, are positively correlated at high significance level across all 31 cancers (Figure 3B). Additionally, miR-483 was found to co-express with all IGF2 isoforms, including INS-IGF2 (insulin, isoform 2) and IGF2-AS (IGF2 antisense RNA). The positive correlation is retained in the normal tissue samples, albeit with compromised significance level due to small sample size. As a control, miR-483 shows little correlation with the housekeeping gene GAPDH in all cancers studied.

In another scenario where multiple miRNAs reside within the same gene, we observed two distinct phenotypes. The miRNA-hosting gene GABRE (Gamma-Aminobutyric Acid Type A Receptor Epsilon Subunit), which encodes a ligand-gated ionic channel, gamma-aminobutyric acid (GABA), significantly positively correlates with both of its intragenic miRNAs, miR-224 and miR-452 in most cancers (Figure 3C). On the other hand, neither of the two embedded miRNAs, miR-1306 and miR-3618, showed significant correlation with their shared host gene DGCR8 (DiGeorge Syndrome Critical Region Gene 8), in any of the studied cancers (Figure 3D). Not surprisingly, DGCR8 as an essential microprocessor complex component is constantly highly expressed across all cancers studied, and exemplifies negligible variance among patient samples (Figure 3E). In contrast, its hosted miRNAs, miRNA-1306 and miR-3618, both generally expressed at low levels in all cancers, and showed much higher inter-sample variation, leading to little correlation with their host gene DGCR8. Additional examples phenocopied the second scenario largely due to negligible expression level of associated miRNAs (Figure S6B-E). Further analysis excluded the potential impact of genomic location of the intragenic miRNA on the positive correlation with its host gene (Figure S6A).

### miRNAs can activate a gene by inhibiting its upstream suppressors

An observed positive miRNA-gene correlation might also result from indirect regulation. Specifically, a miRNA can target and inhibit an intermediate gene, which in turn serves as an upstream suppressor of a gene positively correlated with this miRNA, forming a double-negative loop. To figure out how many positive pairs were governed by this mechanism, we first identified all intermediate (IM) genes for each positive pair that are significantly negatively correlated with both components (the miRNA and gene) of the pair (*R* < −0.1, adj.P < 0.05). To ensure consistency, we required that this double-negative correlation holds significant in at least five cancer types. Under this criterion, we detected no intermediate genes in a large portion of the positive pairs, with percentage of non-IM (*N* = 0) ranging from 70.8% in PanCan10 to 56.9% in PanCan20 groups, respectively (Figure 4A-B, top). When we narrowed down the general IM genes to high-confident miRNA targets (IM_TAR) of the associated miRNA (predicted by TargetScanHuman with high confidence), the percentage further increased to 90.7% in PanCan15 and 83.7% in PanCan20 groups, respectively (Figure 4A-B, bottom). These results indicate that an overwhelming part of the top-ranked positive correlations cannot be explained by this double-negative indirect regulation mechanism (Figure 4C). Interestingly, pairs in high pan-level groups (PanCan20/25) were more likely to bear intermediate genes compared to their counterparts in low pan-level groups (PanCan10/15), in both IM and IM_TAR scenarios.

**Figure 4.**
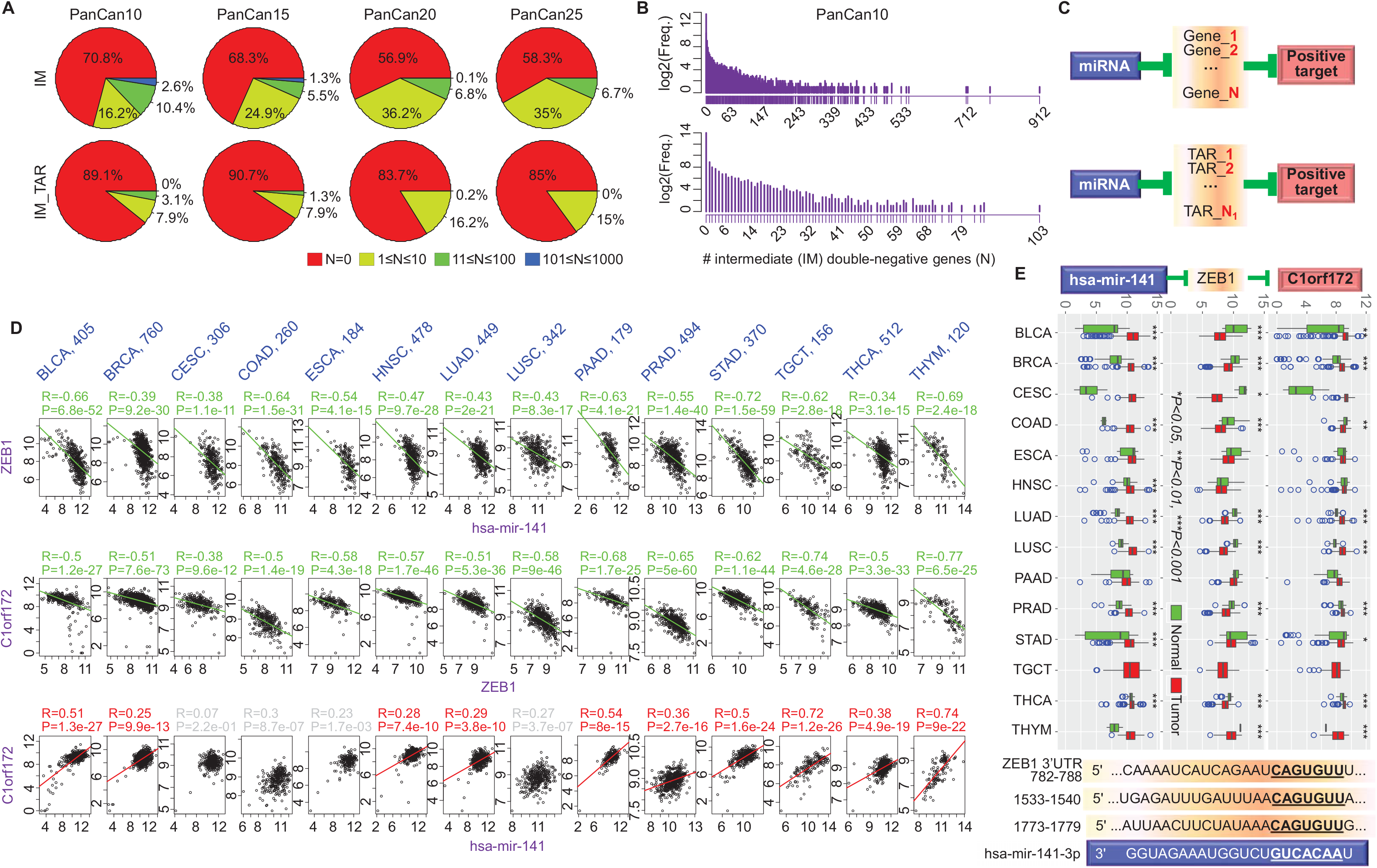
miRNAs can activate a gene by inhibiting its upstream suppressors. (A) Top: Percentage of positive pairs with *N* (0≤*N*≤1000) detected intermediate genes (IM) that are negatively correlated with both (double-negative) the miRNA and gene in the pair at different pan-levels. Bottom: Percentage of positive pairs with *N* (0≤*N*≤1000) IMs predicted to be targets of the associated miRNA in TargetScanHuman (IM_TAR). The number (*N*) of intermediate genes for each pair is color-coded in the legend. (B) Frequency (log2 transformed) distribution of number of intermediate double-negative genes for PanCan10 positive pairs for IM (top) and IM_TAR (bottom) scenarios as in (A). (C) Schematic illustration of double-negative regulation for IM (top) and IM_TAR (bottom) scenarios. (D) A representative example (hsa-mir-141~ZEB1~C1orf172) of the double-negative regulation mechanism for positive correlation (IM_TAR scenario). Top: Correlation between hsa-mir-141 and one of its target genes ZEB1 (predicted by TargetScanHuman with high confidence). Middle: Correlation between ZEB1 and C1orf172, one of those genes positively correlated with hsa-mir-141. Bottom: Correlation between hsa-mir-141 and C1orf172. Correlations were assessed by Pearson correlation coefficient and p-value across 14 TCGA cancers, with sample sizes indicated above corresponding columns. Green: significant negative correlation; Red: significant positive correlation; Gray: non-significant correlation. (E) Top: Expression level of hsa-mir-141, ZEB1 and C1orf172 in the same 14 cancer tissues as in (D), and in comparison with their normal controls if available. The middle line in the box is the median, the bottom and top of the box are the first and third quartiles, the whiskers extend to 1.5 IQR of the lower quartile and the upper quartile respectively, and the blue hollow circles represents outliers. Expression difference significance (*p*-value) was obtained from two-sided student *t*-test. Bottom: Presence of binding sites of hsa-mir-141 on the 3’ untranslated region (3’UTR) of ZEB1 predicted by TargetScanHuman. See also Figure S7.

As a typical example, we detected 912 IM genes and 86 IM_TAR intermediate genes respectively for the positive correlation pair miR-141~C1orf172. Among the 86 IM_TAR genes, a transcription factor gene ZEB1 (zinc finger E-box binding homeobox 1) was found to negatively correlate with both miR-141 and C1orf172 across 14 cancers (Figure 4D). The miR-141~C1orf172 itself was significantly positively correlated in 10 out of the 14 cancers, while in the remaining four cancers the significance is marginal. We further checked the specific expression profiles of these two components in this double-negative loop in the same 14 TCGA cancer tissues. Results showed that miR-141 and gene C1orf172 were highly expressed while the IM_TAR gene ZEB1 downregulated in tumor samples than in normal controls, and the difference was statistically significant in most cancers studied (Figure 4E, top). The correlation and differential expression analysis corroborated that the inhibition effect of miR-141 on ZEB1 was potent, assumedly owing to the three highly conserved binding sites of miR-141 on the 3’UTR of ZEB1 (Figure 4E, bottom). Additional examples regarding positive pairs miR-200c~C1orf172 and miR-429~C1orf172 further confirmed this double-negative regulation mechanism (Figure S7). Interestingly, ZEB1 was largely recognized as an oncogene due to its contribution to epithelial-to-mesenchymal transition (EMT), and hence the miR-141/200 family members were considered as tumor suppressors because they target and potently inhibit ZEB1 expression (Cursons et al., 2018;Gregory et al., 2008). Its general downregulation in tumor samples in various TCGA cancer types further indicates the complexity of gene regulation.

### Positive miRNA-gene correlations can result from co-transcription mechanisms

Analogous to the double-negative regulation, an observed positive miRNA-gene correlation can also result from double-positive correlation. Specifically, a third gene may serve as a transcriptional activator for both the constituent miRNA and gene of a studied pair. To determine how many positive pairs were governed by this mechanism, we first identified all intermediate (IM) genes for each positive pair that are significantly positively correlated with both the constituent miRNA and gene (*R* > 0.1, adj.P < 0.05). To ensure consistency, we required that this double-positive correlation holds significant in at least five cancer types, as in the double-negative scenario. Under this criterion, we found only a negligible portion of pairs having no intermediate genes, with percentage of non-IM (*N* = 0) ranging from 0.9% in PanCan10 to 2.1% in PanCan25 groups (Figure 5A, top). Surprisingly, a considerable portion of pairs presented more than 1000 intermediate genes in all pan-level groups, ranging from 17.4% in PanCan10 to 82% in PanCan20 groups. These patterns are largely distinct from those observed in the double-negative scenario. When we narrowed down the general intermediate genes to general transcription factors (IM_gTF), i.e., those have been reported as a transcription factor in at least one of six previous studies (see Methods), the number of intermediate genes decreased dramatically. The *N* > 1000 group disappeared, while the 101 ≤ *N* ≤ 1000 group replaced it to dominate, and the *N* = 0 group increased a little (Figure 5A, middle). When we further narrowed down the IM_gTF genes to those TFs that have been reported as a specific TF (IM_sTF) of the constituent gene in the pair, the number of intermediate genes further significantly decreased. About 20-30% of pairs lost all intermediate genes, while all the remaining pairs presented less than 100 intermediate genes (Figure 5A, bottom). In contrast to the double-negative scenario, which showed monotonically decreasing number of positive pairs over number of intermediate gene *N*, the number of positive pairs in double-positive scenario displayed a normal-like distribution over number of intermediate genes, especially in the IM and IM_gTF cases (Figure 5B). These results indicate that a large part (70-80%) of the top-ranked positive correlations might be explained by this double-positive co-transcription mechanism (Figure 5C).

**Figure 5.**
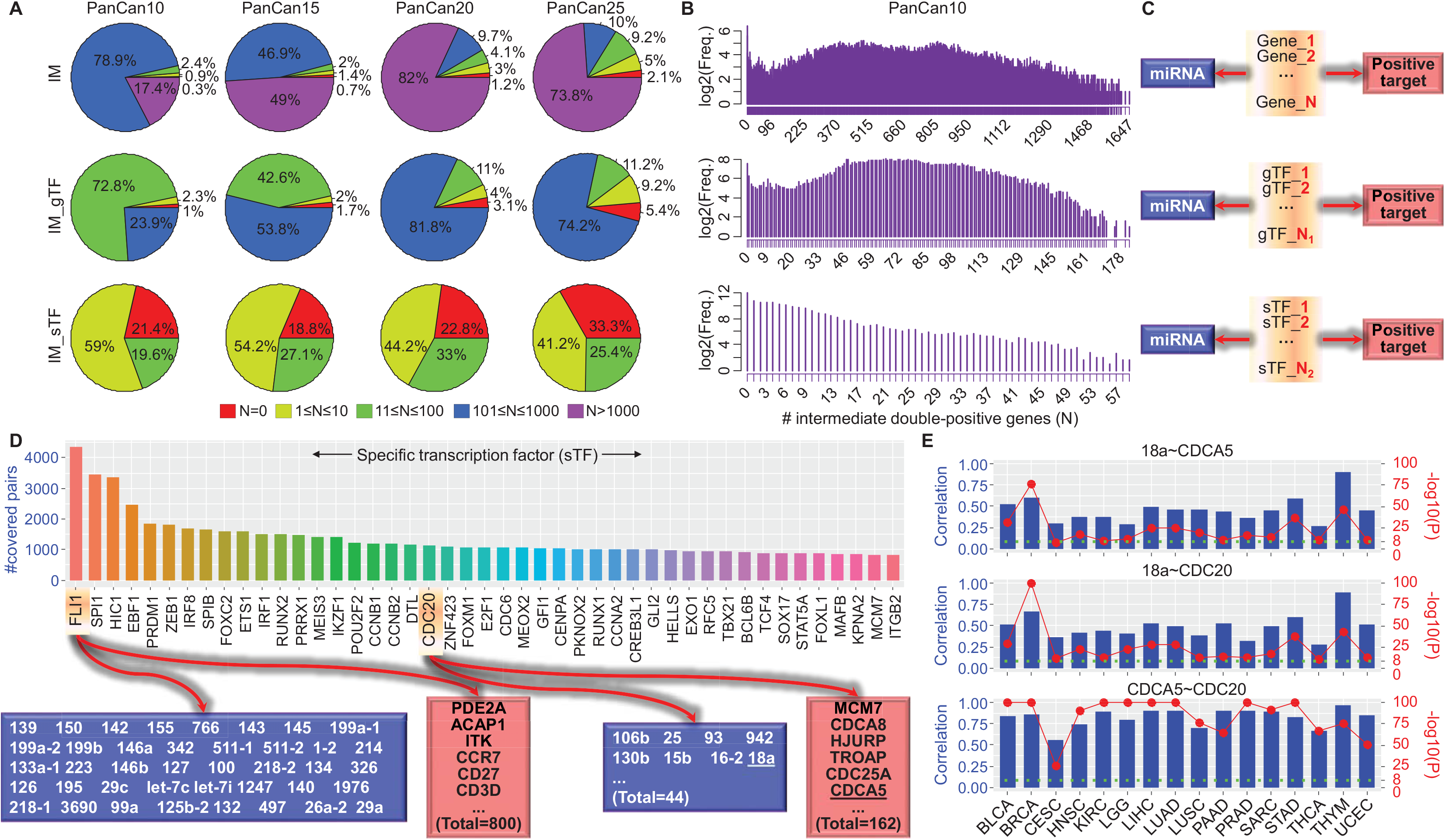
Positive miRNA-gene correlations can result from co-transcription mechanisms.(A) Top: Percentage of positive pairs with *N* (*N*≥0) detected intermediate genes (IM) that are positively correlated with both (double-positive) the miRNA and gene in the pair at different pan-levels. Middle: Percentage of positive pairs with *N* (*N*≥0) IMs previously reported as a general transcription factor (IM_gTF). Bottom: Percentage of positive pairs with *N* (*N*≥0) IMs previously reported as a specific transcription factor targeting on the gene in the positive pair (IM_sTF). The number (*N*) of intermediate genes for each pair is color-coded in the legend. (B) Frequency (log2 transformed) distribution of number of intermediate double-positive genes for PanCan10 positive pairs for IM (top), IM_gTF (middle) and IM_sTF (bottom). (C) Schematic illustration of the double-positive regulation (co-transcription) for IM (top), IM_gTF (middle) and IM_sTF (bottom) scenarios. (D) Specific transcription factors (sTF) appearing in multiple positive pairs. The sTFs are ranked by the number of positive pairs they covered plot (top), with two representative examples for TFs FLI1 (involved in 4357 pairs) and CDC20 (involved in 1145 pairs) detailed (bottom). (E) A representative example showing transcription factor CDC20 was significantly positively correlated with both the miRNA and gene in a pair across 16 TCGA cancer types. Pairwise correlation among the miRNA hsa-mir-18a, the gene CDCA5 and the associated TF CDC20 are shown. Pearson correlation coefficient and significance level (-log10 transformed) are illustrated as blue bars and red solid circles respectively. A threshold -log10(P)=8 is marked by a green dotted line. See also Figure S8.

We then focused on the specific transcription factors that served as the intermediate double-positive genes (IM_sTF). We noted that TFs tended to be shared by multiple positive pairs, and each of the top 45 specific TFs covered at least 800 pairs (Figure 5D, top), implying that some transcription factors are highly active and tend to function as common activators of many target genes. Striking examples include the friend leukemia integration 1 (FLI1) transcription factor, which presented as an IM_sTF of 4357 pairs spanning 40 miRNAs and 800 genes, and the cell division cycle protein 20 (CDC20), which appeared in 1145 positive pairs involving 44 miRNAs and 162 genes (Figure 5D, bottom). Looking into the correlation details regarding the detected double-positive loop miR-18a~CDC20~CDCA5, we found that they were positively correlated in 16 TCGA cancers in a pairwise manner (Figure 5E). FLI1 has been recognized as a prototype oncogene, whose aberrant expression is associated with chromosome abnormalities leading to an oncogenic fusion gene EWSR1/FLI1 with strong transforming capabilities (Ohno et al., 1993;Rao et al., 1993). Additional examples regarding FLI1’s correlation in miR-139~PDE2A and miR-142~ITK pairs further confirmed its critical role in gene regulation (Figure S8). CDC20 is a regulatory factor interacting with other cell cycle related proteins and has recently been claimed as a potential novel therapeutic target for cancer treatment (Wang et al., 2013). These results documented the biological significance of the detected intermediate genes as specific transcription activators, and consolidated the potential co-transcription mechanism.

### miRNAs/genes with positive correlations tend to form super-enhancer-like regions

Super-enhancers are an emerging subclass of regulatory regions that have been suggested to drive the processing and production of master miRNAs crucial for cell identity and oncogenesis (i.e. associated with cancer hallmarks) (Suzuki et al., 2017). Super-enhancers are described as a type of enhancer domains that are densely occupied by the master regulators (Oct4, Sox2 and Nanog) and Mediator (Whyte et al., 2013). In the established ROSE (Rank Ordering of Super-Enhancers) pipeline (Hnisz et al., 2013), super-enhancers were identified based on the ChIP-seq signal of a typical enhancer marker histone H3 lysine 27 acetylation (H3K27ac). We analyzed the H3K27ac ChIP-seq data of 64 samples across 15 human tissues from ENCODE, based on a modified ROSE pipeline (see Methods). We detected significant enrichment of H3K27ac signal surrounding the miRNA/gene sites associated with the top-ranked positive pairs. The top 64 miRNAs ranked by their average ROSE score over 15 tissues displayed discernible patterns in different tissue types (Figure 6A). The top 64 miRNAs themselves can also be clustered into several groups with clear group-specific signatures. These results verified previous findings that super-enhancers are able to mark master miRNAs crucial for cell identity.

**Figure 6.**
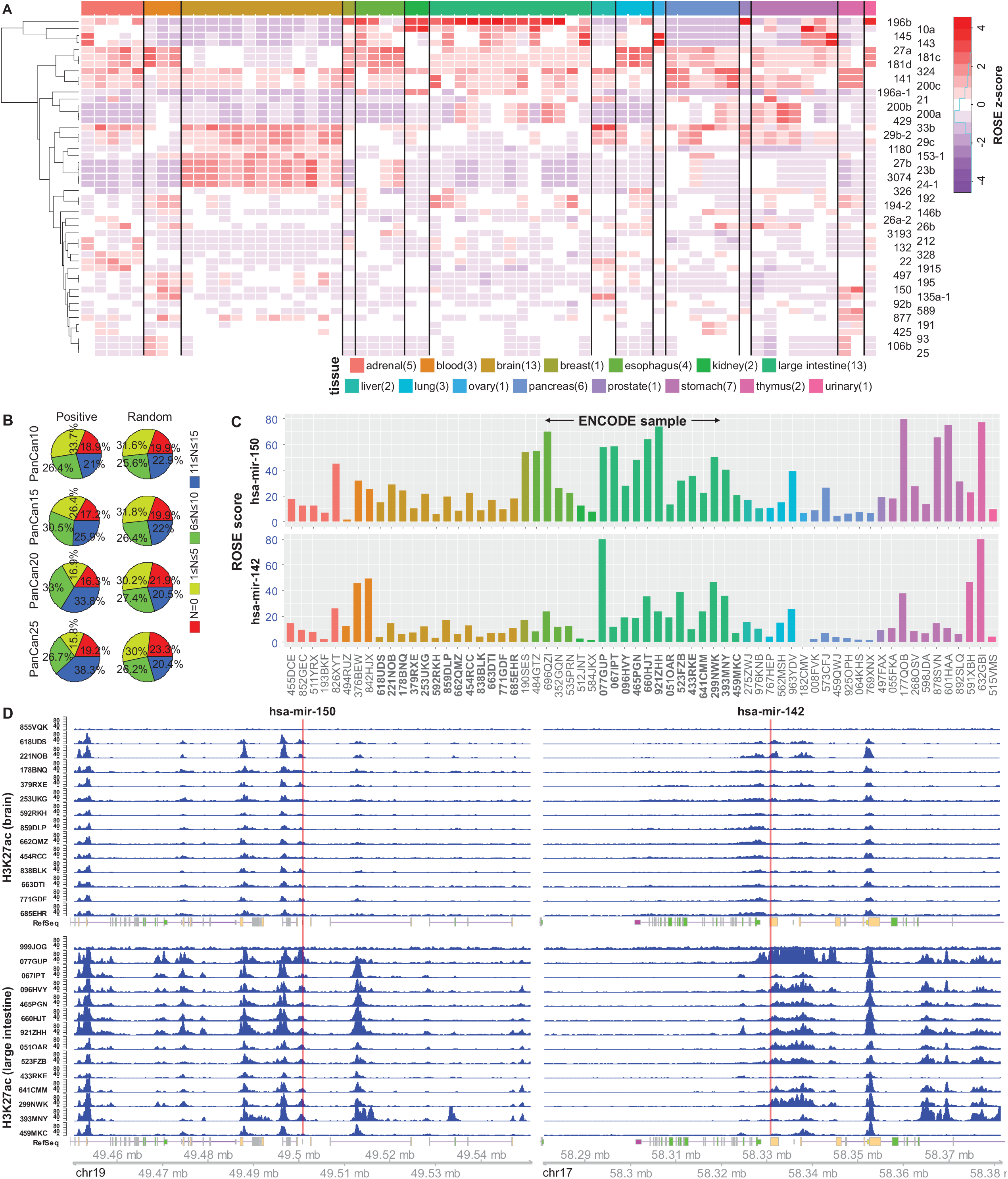
Positive correlation associated miRNAs/genes tend to form super-enhancers. (A) Relative ROSE score of H3K27ac signal surrounding sites of the top 64 miRNAs from top positive pairs covering ≥10 cancers (PanCan10). ROSE scores were centered and scaled across miRNAs for each ENCODE sample before clustering over miRNAs. Names of 64 miRNAs and 15 ENCODE tissues are shown on the right and at the bottom of the heat map, respectively. Color-key for normalized ROSE score is shown on the top right corner. (B) Percentage of significantly positive (Positive, left) or randomly chosen (Random, right) miRNA-gene pairs with high ROSE score (≥10) on either the miRNA or gene site in each pair at different pan-levels. The Random group consists of same number of miRNA-gene pairs as the Positive group at each pan-level, with constituent miRNAs and genes randomly chosen from those not involved in the PanCan10 pairs. The number (*N*, 0≤N≤15) of ENCODE tissues covered by a pair is shown in the legend. (C) A representative example showing ROSE score of H3K27ac signal for two miRNAs hsa-mir-150 (top) and hsa-mir-142 (bottom) in 64 ENCODE samples across 15 human tissues. The color code for each tissue type is same with (A). (D) Detailed illustration of H3K27ac signal intensity surrounding ±50K bp of hsa-mir-150 (left) and hsa-mir-142 in ENCODE brain (top) and large intestine (bottom) tissues, respectively. Genomic coordinates for each region is shown at the bottom. The TSS of each miRNA is marked by a red solid line. Peak regions were binned to windows of 200bp for visualization of the H3K27ac ChIP-seq signal. See also Figure S9-S10 and Table S5 for more results of ROSE analysis on miRNA/gene sites.

We then compared the H3K27ac enrichment profiles between positive pairs and pairs randomly chosen from the complementary set of PanCan10 group. The percentage of pairs covering more than 10 out of 15 ENCODE (11≤*N*≤15) tissues steadily increased with pan-levels (from 21% in PanCan10 to 38.3% in PanCan25) in the positive pair group, while the randomly chosen pairs remained at much lower percentage across pan-levels (Figure 6B). Frequency distribution analysis of the ROSE score of the PanCan25 positive pairs and randomly chosen pairs showed a right shift of the PanCan25 group (Figure S9), indicating that positive pairs tend to yield a higher ROSE score than random ones. We then went into the detail of two representative miRNAs miR-150 and miR-142, which ranked the top 1st and 3rd based on number of positive targets (Figure 1B) respectively, and detected high intensity of H3K27ac signal surrounding their TSS sites across all 15 ENCODE tissues studied (Figure 6C). Furthermore, while inter-tissue variation exists, samples of same tissue type showed considerable consistence in H3K27ac enrichment at the miR-150 and miR-142 loci, as exemplified by the two representative tissues brain and large intestine (Figure 6D). By checking the top 10 pairs enriching for H3K27ac signal, we found miR-196b/150/142/155 and the HOX family genes are involved most frequently (Figure S9-10). Together, these results reveal that the positive correlation associated miRNA/gene sites are more likely to enrich H3K27ac histone modification in a special manner that mimics the established super-enhancer pattern.

### Positive miRNA-gene correlations are partially retained in *in vitro* conditions

To check whether the positive miRNA-gene correlations detected in patient samples are also retained in the *in vitro* condition, we conducted a series of wet-lab experiments in several cancer cell lines as well as BJ primary human fibroblast cells. We selected as prototype 8 miRNA-gene pairs that are discernably detectable and positively correlated across TCGA cancers, including miR-21~TGFBI, miR-142~CTLA4/IL7R/PIK3CG, miR-155~CTLA4/ITK, and miR-214~PDGFRA/IGFBP5 (Figure S11A-B). Among these 8 pairs, miR-21~TGFBI and miR-214~IGFBP5 are not in the PanCan10 group. All of the 8 pairs except miR-214~PDGFRA possess miRNA binding sites in the 3’UTR of the corresponding partner gene (Figure S11C). To select proper cell lines for experiments, we checked the gene expression in several candidate cell lines derived from human cancers, including breast cancer (MDA-MB-231 and T47D), liver cancer (Huh7) and cervical cancer (HeLa), as well as primary human fibroblast cells (BJ). We found that these genes are expressed in a highly cell line-specific manner (Figure 7A, bottom and Figure S11D). However, these genes are generally highly expressed in all TCGA cancer types (Figure 7A, top and Figure S11B), which demonstrates difference between *in vitro* and *in vivo* conditions.

**Figure 7.**
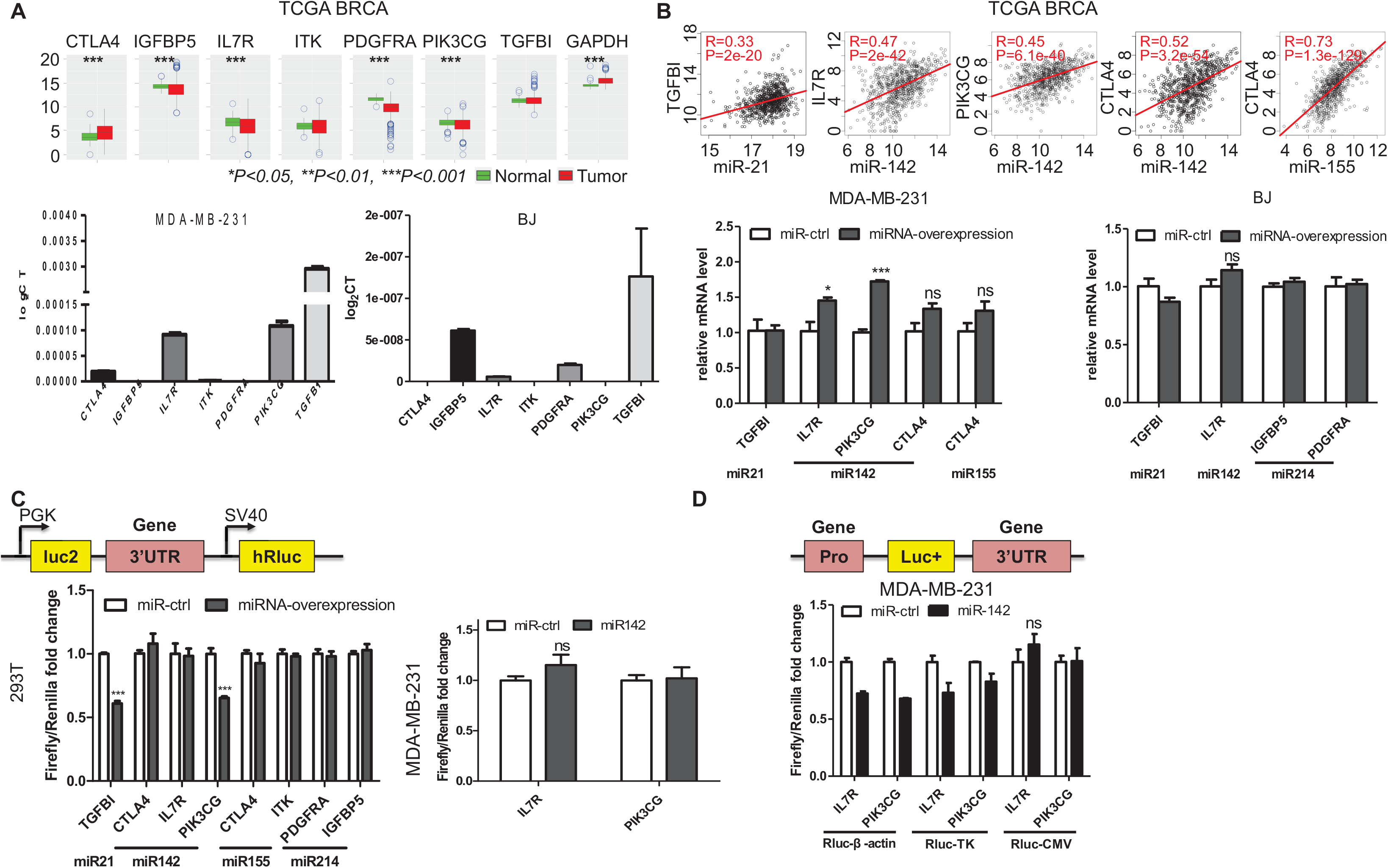
The positive regulation of gene expression by miRNAs detected in patient samples is partially recapitulated in cell lines. (A) Top: Boxplot illustrating gene expression in TCGA BRCA cancer samples and associated normal controls for genes in the selected 8 miRNA-gene pairs. Housekeeping gene GAPDH is also shown for control. Comparison was performed by two-sided student *t*-test. Color and shape code for boxplot is as in Figure 4E. Bottom: Endogenous expression level of the same genes, detected by qRT-PCR, in MDA-MB-231 and BJ cell lines as indicated. (B) Top: Pearson correlation between indicated miRNA-gene pairs in TCGA BRCA cancer samples. These pairs are selected since only these associated genes were endogenously expressed at detectable levels in the MDA-MB-231 breast cancer cell line. Bottom: qPCR results showing endogenous gene expression levels with or without stable overexpression of the constituent miRNAs in the selected positive pairs in MDA-MB-231 cells (left) and BJ cells (right). (C) Left: 3’UTR reporter assays showing the activity of the reporters with or without miRNA overexpression for the 8 selected positive pairs in 293T cells. Right: 3’UTR reporter assays showing the activity of the 3’UTR reporters of the IL7R and PIK3CG genes with or without miR-142 in MDA-MB-231 cells. Correlations of these pairs in all TCGA cancers are illustrated in Figure S11A. There is no miR-214 binding sites on its partner gene PDGFRA; for the other 7 pairs, there is at least one conserved miRNA binding site on its partner gene, see Figure S11C. (D) Dual-luciferase assay results of reporters containing both a 3-kb promoter and the3’UTR carrying the miR-142 binding sites of the IL7R or PIK3CG genes in MDA-MB-231 cells. The promoter-3’UTR Firefly luciferase reporters were cotransfected into MDA-MB-231 cells with Renilla Luciferase and corresponding miR-142 or vector control (Ctrl). Luciferase activity was determined 48 hours after transfection. Firefly luciferase activity was normalized to the Renilla luciferase activity. The experiment was done for 3 times, with β-actin, TK, and CMV driven renilla, respectively, and result was displayed separately. In all wet-lab experiments, values are mean ± SD for triplicates. See also Figure S11. The experiment is done for 3 times, 1 with b-actin drived renilla, one with TK drived renilla and one with CMV-renilla, in every single experiment, triplicate wells of cells are collected. This figure conbine the 3 times result together

We further found that the positive regulation was retained in only two pairs miRNA-142 ~ (IL7R, PIK3CG) in MDA-MB-231 breast cancer cells, while in the other pairs, there was either no positive regulation at all or positive regulations that were consistent but not statistically significant, in the breast cancer or BJ cells (Figure 7B bottom), in contrast to all positive correlations in TCGA samples (Figure 7B top). Then we investigated the mechanisms underlying this positive regulation of IL7R and PIK3CG by miR-142 in the subsequent experiments. We did 3’UTR reporter assay in two cell lines, 293T (Figure 7C left) and MDA-MB-231 (Figure 7C right), and found that co-expression of miR-142 did not increase the 3’UTR reporters of IL7R and PIK3CG, suggesting that the positive regulation is not mediated by 3’UTR, which is different from the mechanism for the negative regulation of genes by miRNA. Next, we combined promoter and 3’UTR of each gene in the reporter, and miRNA still has no positive effects on these reporters (Figure 7D), implying that the regulation is not achieved via the promoters or the combination of the promoters and 3’UTRs. Taken these results together, we conclude that the positive regulations observed in tumors are only partially retained in cancer cell cultures, and that the mechanism underlying the positive regulation *in vitro* is not mediated via promoters or 3’UTRs of the genes, even though the promoters and/or 3’UTRs contain the miRNA binding sites. It is possible that the mechanisms for the positive regulations of genes by miRNAs *in vitro* differ from those *in vivo.* Alternatively, the positive correlation might be mediated by certain epigenetic modifications, which cannot be detected on transiently transfected reporters that are not integrated into the original locus of the genes.

## DISCUSSIONS

While miRNAs have recently been reported to activate genes in particular contexts based on *in vitro* cell culture experiments, the phenomenon and its clinical relevance have not been established in *in vivo* conditions. By a comprehensive investigation on the expression correlation between all 1046 miRNAs and 20531 genes in 8375 patient samples across 31 major human cancers in The Cancer Genome Atlas (TCGA), we detected compelling positive miRNA-gene correlation patterns that are highly consistent (conserved) across cancer types. We further confirmed that these conserved positive correlations bear biological significance and clinical relevance. In addition, we identified potential mechanisms underlying the upregulation of genes by miRNAs, including the intragenic miRNA-host gene co-expression, miRNA inhibition of the upstream suppressor of the gene (double-negative), co-transcription by shared transcription factors (double-positive), super-enhancer mediated miRNA-gene co-expression, and direct binding of miRNA to the regulatory regions of the partner gene (Figure 8). Therefore, the present study established a striking phenomenon regarding miRNA-directed gene activation in human cancer samples, confirmed its biological and clinical significance, and partially explained the potential mechanisms underlying this activation, which deserve further experimental exploration.

**Figure 8.**
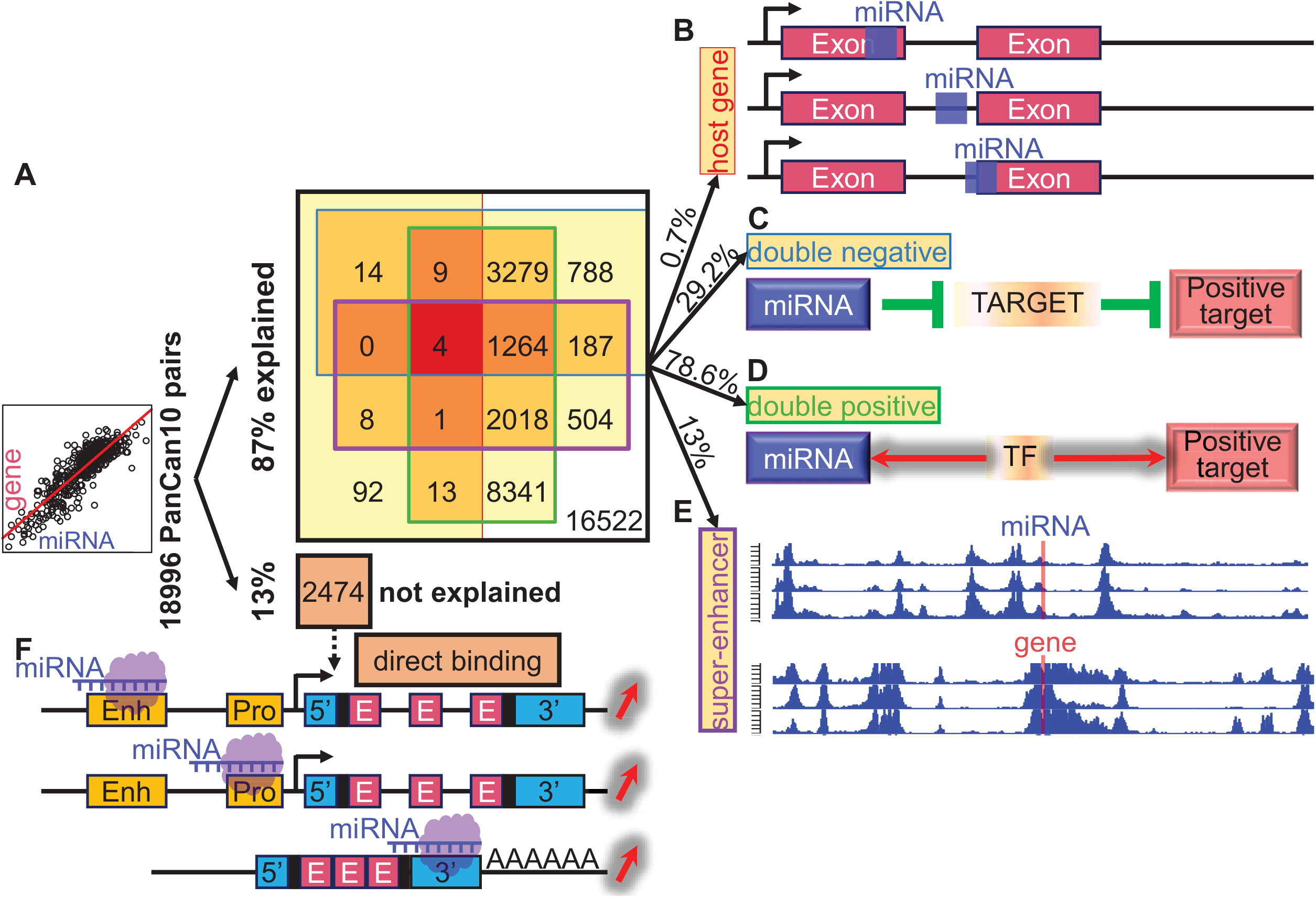
A summary of putative mechanisms underlying miRNA-mediated gene activation. (A) Numbers and percentages of positive pairs (PanCan10, n=18996) explained by various potential mechanisms. (B) Schematic illustration of intragenic (exonic, intronic, and mixed) miRNA and its host gene. (C) Schematic illustration of how miRNAs activate a gene by inhibiting its upstream suppressor. (D) Schematic showing miRNA and gene are positively correlated due to co-transcription by shared transcription factor (TF). (E) Schematic depicting either the miRNA or gene in a pair is enriched for super-enhancer marker H3K27ac. (F) Schematic illustration of how miRNAs activate a gene by directly binding to gene on its enhancer (Enh), promoter (Pro), or 3’UTR region. See also Table S6.

Besides revelation of these unusual and seemingly counterintuitive profiles of miRNA-gene correlations, we also confirmed their biological and clinical significance through a series of stringent analysis. The constituent miRNAs in our detected PanCan10 positive pairs covered all previously reported cancer hallmarks miRNAs, while the constituent genes were significantly enriched in the established 50 hallmark processes pertaining to cell division and proliferation, demonstrating their implications in oncogenesis. Moreover, with increasing pan-levels, the constituent genes converge into a smaller set of biological signaling pathways and cellular processes regarding immune cell differentiation and membrane signaling cascades, indicating that the positive correlations largely participate in the immune response and extracellular message stimulation and subsequent intracellular transduction (Figure 2 and Figure S4). Most strikingly, both miRNAs and genes across all pan-levels exhibit dramatically higher capability in stratifying patients in terms of overall survival (OS) rate, compared to the remaining miRNAs/genes. In most cases, combined miRNA and gene stratifiers perform better than individual ones (Figure 2 and Figure S5). These findings argue that the miRNAs and genes with positive correlation and the positive correlations themselves are clinically significant, rather than random events.

Although intragenic miRNAs tend to positively correlate with their host genes in expression as expected, we found a considerable number of counterexamples (Figure 3D-E and Figure S6). Our further analysis verified the previous hypothesis that conserved miRNAs (conservation adopted from TargetScanHuman v7.2, see Methods) are more likely to co-express with their host genes. However, this hypothesis still could not account for all observations. We checked whether the relative location of a miRNA in its host gene plays an import role in determining their co-expression but found little association. We proposed two possible reasons for the exceptions. First, a house keeping or likewise gene tends not to show any correlation with its embedding miRNAs, since this type of genes are critical for cell vitality and always express at high levels. Examples include DGCR8, which is an essential member of the microprocessor machinery complex and shows little correlation with its intragenic miRNAs miR-3618/1306 (Figure 3D); and CDC37, an important partner of HSP90 participating the regulation of protein kinases, displays poor correlation with its intragenic miRNA miR-1181 (Figure S6C). Second, some miRNAs do not express at all or express at very low level across all cancer tissues, and hence exemplify little if any correlation with their host genes. Examples include miR-1182~ FAM89A and miR-1183~SP4 (Figure S6D-E). These miRNAs might be just putatively derived from conserved hairpin sequences, rather than well-established and well-annotated miRNAs. Another possibility is that there is additional layer of regulation at the level of miRNA biogenesis.

A straightforward question pertaining to the miRNA-gene positive correlation is whether this correlation is direct or indirect; and a more fundamental question is whether this correlation reflects real causation or is just an association of two unrelated events in a statistics sense. We found that most of the observed positive correlations can be well explained by the indirect regulation mechanism, including double-negative (29.2%) and/or double-positive (78.6%) regulation routines (Figure 4-5, 8). In the double-negative scenario, a miRNA inhibits its target genes by direct binding to their 3’UTR, and those target genes in turn suppress their downstream genes in a same signaling pathway. This will lead to an observation of positive correlation between the miRNA and these downstream genes. In addition, such double-negative regulation might be repeated multiple times in a pathway resulting in multiple intermediate layers between the miRNA and its positively correlated genes. While such type of correlation is indirect, it is causative. In the double-positive scenario, on the other hand, components of a positively correlated pair are regulated by shared regulatory factors and seem not causally linked. However, this type of positive correlation has also been reported to be implicated in tumorigenesis (Lim et al., 2013). This is sensible since not all miRNA-gene pairs sharing regulatory factors exemplify positive correlation due to many complex regulation networks, and cancer cells take advantage of this switch to control miRNA-gene co-expression. For example, a genetic variation in a regulatory region can impact the binding affinity of miRNAs or transcription factors, leading to abnormal gene abundance (Tan, 2017, 2018). Notably, the other two potential mechanisms, miRNA-host gene co-expression, and super-enhancer mediated miRNA-gene co-expression, might also be ascribed to a mechanism similar to the dual-positive regulation. Striking examples include the FLI1 regulated miR-139 and PDE2A co-upregulation (Figure S8B), and H3K27ac enrichment associated miR-196b and HOXA10 co-activation (Figure S9-10). We also noticed a significant overlap between different potential mechanisms as elucidated above (Figure 8). In particular, four pairs, including 152~COPZ2, 106b~MCM7, 25~MCM7 and 93~MCM7, can be explained by either of the four proposed mechanisms (Table S6). We conclude that in a more general sense, all of the four hypotheses (Figure 8B-E) are indirect regulation that accumulatively accounts for the majority (87%) of the PanCan10 positive pairs.

The more exciting scenario for miRNA-directed gene upregulation is through direct binding and activation. Previous studies confirmed that miRNA can activate a gene by direct binding to its regulatory elements at the RNA or DNA level including 3’UTR (Vasudevan et al., 2007), 5’UTR (Liu et al., 2013), promoter (Huang et al., 2012;Majid et al., 2010;Place et al., 2008;Zhang et al., 2014), or enhancer (Xiao et al., 2017) regions, as abovementioned. In our study, a small portion of miRNA-gene positive correlations (13%) that could not be accounted for by any of our proposed four indirect regulation hypotheses might be explained by the direct binding and activation hypothesis. It should be realized that this hypothesis necessitates further experimental validation. Actually, even the indirect regulation hypotheses as discussed above only provide candidate intermediate regulators to be narrowed down by elaborately designed experiments. Our *in vitro* experiments partially verified that miR-142 overexpression can activate IL7R and PIK3CG in human breast cancer cells line MDA-MB-231 (Figure 7B). However, in other cases, our wet-lab experiment results showed distinct miRNA-gene interaction patterns in comparison with those observed in corresponding TCGA cancer samples. This discrepancy argues the importance of the critical impact of various *in vivo* microenvironment factors, e.g., the intercellular communications and the immune responses, on the miRNA-gene interactions. And this argument is further consolidated by our enrichment analysis revealing that cell cycle related processes, cell membrane signaling and immune cell differentiation are involved in the top-ranked positive pairs (Figure 2). Therefore, our comprehensive analysis based on large sets of patient samples is essential for the revelation of phenomenon, biological significance and potential mechanisms underlying the miRNA-mediated gene activation.

## MATERIALS AND METHODS

### TCGA data acquisition, quality control and preprocessing

The microRNA (miRNA) and gene expression data, as well as clinical information of each cancer, were downloaded from the Cancer Genome Atlas (TCGA) data portal (https://portal.gdc.cancer.gov/). Three cancers, including FPPP (FFPE Pilot Phase II), GBM (glioblastoma multiforme) and LAML (acute myeloid leukemia) were excluded due to small sample size or platform inconsistency. The pan-cancer analyses eventually consisted of 31 cancer types: ACC, BLCA, BRCA, CESC, CHOL, COAD, DLBC, ESCA, HNSC, KICH, KIRC, KIRP, LGG, LIHC, LUAD, LUSC, MESO, OV, PAAD, PCPG, PRAD, READ, SARC, SKCM, STAD, TGCT, THCA, THYM, UCEC, UCS and UVM. Sample size ranges from 36 (CHOL) to 1102 (BRCA), see Table S1. A total of 1046 miRNAs and 20531 genes (protein coding and noncoding) were included in the TCGA IlluminaHiseq miRNASeq and IlluminaHiSeq RNASeqV2 data, respectively. The miRNA and gene expression values were logarithmically transformed (base 2) prior to further analysis. Pearson correlation analysis was performed to assess the co-expression between miRNAs and genes across all 31 cancer types. A correlation was deemed significant in a cancer if its absolute correlation coefficient |*R*| > 0.1 (R > 0.1 for positive correlation and *R* < −0.1 for negative correlation) and Hochberg adjusted *p*-value adj.P < 0.05 unless otherwise stated.

### Functional characterization of positive correlation related miRNAs and genes

To explore the biological significance of the top-ranked pairs, we conducted gene ontology (GO) and KEGG (Kanehisa et al., 2017) signaling pathway enrichment analyses on the genes involved in the top-ranked significant pairs, using the R package clusterProfiler (Yu et al., 2012).

Enrichment profiles were checked over genes at different pan-levels (PanCan10/15/20/25) respectively, with Fisher exact test p-value < 0.05 deemed significant. To further investigate the clinical relevance of these pairs, we performed survival analysis on different groups based on their miRNAs and gene expression profiles. Briefly, patients were first divided into non-overlapped groups based on their miRNA and gene expression in three ways: 1) patients were divided into three groups as high-middle-low (Hi-Mi-Lo) by miRNA expression; 2) patients were divided into three groups as high-middle-low (Hi-Mi-Lo) by gene expression and 3) patients were divided into two groups as high-low (Hi-Lo) by gene:miRNA ratio. Then we adopted log-rank test to compare the overall survival probability of patients from different groups, categorized by single or combined stratification indexes. Therefore, for a particular miRNA~gene pair, the HiHi+Hi group refers to patients from high miRNA expression, high gene expression and high gene:miRNA ratio group. The stratification power in patient survival of miRNAs/genes from different pan-level groups was compared by the Kolmogorov-Smirnov test (K-S test) performed on the density distribution of their log-rank test p-values.

To further investigate the implications of our detected positive correlations in tumorigenesis, we associated the top-ranked miRNA and genes to the well-known cancer hallmark traits. Briefly, we checked the association of the PanCan10 miRNAs with the well-known 10 cancer hallmarks proposed by Hanahan and Weinberg 2011(Hanahan and Weinberg, 2011), as did in a previous study (Suzuki et al., 2017). We also examined the enrichment of the top-ranked genes of different pan-levels in 50 well-established hallmark gene sets related to critical biological processes regarding cellular component, development, DNA damage, immune, metabolic, pathway, proliferation and signaling (Liberzon et al., 2015), which are assumedly essentially relevant to cancer initiation and progression.

### Identification of miRNAs with host genes

We downloaded the gene annotations (hg38) of 27423 genes (including 19902 protein-coding genes and 7521 long non-coding RNAs – lincRNAs) from GENCODE (GTF v25) (Harrow et al., 2012), and annotations of 1881 human miRNAs from miRBase (hsa.gff3) (Kozomara and Griffiths-Jones, 2014). We determined whether a miRNA is embedded in another gene (protein-coding or noncoding RNA) according to their genomic coordinates (locations) from the annotation files. A total of 591 miRNAs were found to be located inside a specific gene, termed “host” gene. Of these, 451 pairs were covered in the TCGA data. These 591 host genes consisted of 474 protein-coding genes and 117 lincRNAs, see Table S4. We investigated the preference of a miRNA to co-express with their host gene by hypergeometric test. The conservation information of each miRNA was adopted from TargetScanHuman v7.2 (miR_Family_Info.txt). A positive conservation score (Conservation? = 1, 2) of a miRNA indicates conservation while a negative score (−1) refers to non-conservation, remaining miRNAs with a zero score were ignored (Table S4).

### Detection of double-negative patterns underlying positive correlation

To validate the hypothesis that a miRNA can upregulate a gene by inhibiting its upstream suppressor, we attempted to detect the double-negative patterns. For a significant positive miRNA-gene pair, we first detected all the intermediate (IM) genes that negatively correlate with both the miRNA and gene in the pair across multiple cancers (*R* < −0.1, adj.P < 0.05, cancer coverage≥5). Then we narrowed down the intermediate genes to real targets of the miRNA based on TargetScanHuman v7.2 (Agarwal et al., 2015). We downloaded the predicted targets (TAR) with context++ and weighted context++ scores, followed by a series of preprocessing steps, including extraction of human species, parsing/trimming miRNA names, removing duplicates, and eventually obtained 198,312 targeting records with high confidence, which involves 321 different miRNAs and 13035 genes.

### Detection of double-positive patterns underlying positive correlation

To validate the hypothesis that a positively correlated miRNA-gene pair might be regulated by shared transcription activators, we tried to detect the double-positive patterns. For a significant positive miRNA-gene pair, we first detected all the intermediate (IM) genes that positively correlate with both the miRNA and gene in the pair across multiple cancers (*R* > 0.1, adj.P < 0.05, cancer coverage ≥ 5). Then we narrowed down the genes by two steps. First, we restricted the IMs into general transcription factors (gTF). We downloaded transcription factors (TF) and their targets with the R data package tftargets (https://github.com/slowkow/tftargets). This dataset includes human TF information curated from six published databases: TRED (Jiang et al., 2007), ITFP (Zheng et al., 2008), ENCODE (Consortium, 2012), Neph2012(Neph et al., 2012), TRRUST (Han et al., 2015), Marbach2016(Marbach et al., 2016). We mapped the Entrez gene IDs into gene symbols using two R packages: annotate (<u>10.18129/B9.bioc.annotate</u>) and org.Hs.eg.db (<u>10.18129/B9.bioc.org.Hs.eg.db</u>). After integration and removal of duplicates, we obtained 2705 gTFs with their target genes, based on which we removed intermediate genes of each pair that were not included in the gTF sets. At the second step, we further narrowed down the gTFs into specific transcription factors (sTF) by removing gTFs whose target genes do not include the gene in the miRNA-gene pair under investigation.

### Super-enhancer (SE) identification from ENCODE H3K27ac ChIP-seq data

We downloaded the H3K27ac ChIP-seq data (bed files for narrowPeak) of 15 human tissues including 64 samples from ENCODE (including blood, lung, liver, kidney, brain, large intestine, stomach, pancreas, esophagus, prostate gland, adrenal gland, breast, ovary, thymus and urinary bladder tissues, see Table S5) (Consortium, 2012). To detect the super-enhancer formation profile, we investigated the H3K27ac signal surrounding a miRNA or gene site based on the ROSE (Rank Ordering of Super-enhancers) pipeline (Loven et al., 2013) with minor modifications. Briefly, we first stitched the detected peaks if they are within a certain distance (called the “region”), and then calculated the average input-subtracted H3K27ac signal intensity within the region by a revised ROSE score:

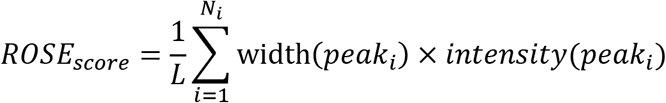

In this formula, *N_i_* is the number of peaks detected in the region, and *L* is the length (in bp) of the region in question. In this study, *L*=100K bp, centered at the transcription start site (TSS) of each miRNA or gene. The width and intensity of each peak (*peak_i_*) was obtained from its genomic coordinates (*width* = chromEnd - chromStart) and signal intensity (signalValue), respectively (see ENCODE narrowPeak format description). We employed the scaled peak intensity (“score” in the bed file) instead of signalValue for better visualization. As did in the original ROSE pipeline, we adopted a promoter exclusion zone of 4K bp, i.e., if a peak was entirely contained within a window of ±2K bp around the TSS, the peak was excluded from the calculation. It should be noted that in contrast to the original ROSE pipeline, we removed a limitation on the maximum distance of 12.5K bp between two constituent enhancers, to focus on the general H3K27ac intensity surrounding the miRNA/gene TSS site. Under our framework, a region with *ROSE_score_*≥10 in at least 11 out of 15 tissues was considered as a super-enhancer (SE).

### Cell culture

BJ human foreskin fibroblasts were maintained in minimum essential medium supplemented with 10% fetal calf serum, nonessential amino acids, and antibiotics. 293T, MDA-MB-231 and Huh7 cells were grown in Dulbecco’s modified Eagle medium supplemented with 10% fetal calf serum, glutamine, and antibiotics. Hela, Huh7 and T47D cells were cultured in EMEM, DMEM and RPMI-1640, respectively, all basic medium was supplemented with 10% FCS and 1% antibiotics.

### Plasmids

The expression vectors for miRNAs and 3’UTR dual-luciferase reporter plasmid (pmirGLO) were purchased from Biosettia, Inc (San Diego, CA). To construct target gene 3’UTR dual-luciferase reporters (pmirGLO-CTLA4-3’UTR, pmirGLO-IGFBP5-3’UTR, pmirGLO-ITK-3’UTR, pmirGLO-PDGFRA-3’UTR, pmirGLO-PIK3CG-3’UTR, pmirGLO-TGFBI-3’UTR, pmirGLO-IL7R-3’UTR), target gene 3’UTR exons containing miRNA seed sequences were amplified by PCR from the genomic DNA of 293T cells. The PCR primers are described in supplemental Materials. All 3’UTR fragments are inserted into pmirGLO by NheI-HF and XhoI.

The pGL3-basic plasmid was purchased from Promega Corporation. To construct pGL3-IL7R/PIK3CG-3Kb+3’UTR reporter, a 3kb promoter sequence from the IL7R and PIK3CG gene was amplified from the genomic DNA of 293T cells and inserted into pGL3-basic plasmid by NheI and XhoI (IL7R promoter), or MluI and BglII (PIK3CG promoter). Then, the PCR product for the 3’UTR of IL7R or PIK3CG (still containing the miRNA seed sequence) was digested by XbaI and inserted into pGL3-IL7R/PIK3CG-3Kb reporter, respectively, downstream of luc+. The PCR primers are described in supplemental Materials.

### Lentivirus-based gene transduction

pLV-miR-ctrl, pLV-miR-21, pLV-miR-142, pLV-miR-155, pLV-miR-214 Recombinant lentiviruses were packaged in 293T cells in the presence of helper plasmids (pMDLg, pRSV-REV, and pVSV-G) using Lipofectamine 2000 (Invitrogen). BJ or MDA-MB-231 cells (1 × 10^5^/well) were seeded into 6-well plates, grown overnight, infected with 300 ul virus in 3 ml fresh medium containing 8 μg/mL polybrene, and spun for 1 hour at 1,600 to 1,800 rpm. Transduced cells were purified with 1.2 μg/mL of puromycin.

### RNA isolation and quantitative Real-Time PCR

RNA was isolated from cells using TRIzol (Thermo Fisher Scientific) according to manufacturer’s protocol. 500 ng of RNA was reverse transcribed to cDNA with iScript^TM^ Reverse Transcription Supermix(Bio-Rad). Quantitative real-time PCR was performed in triplicate with gene-specific primers and SsoAdvance^TM^ SYBR Green Supermix (Bio-Rad) in a Bio-Rad CFX96 REAL TIME SYSTEM following manufacturer’s protocols. GAPDH was used as internal control to normalize the mRNA input for each gene. qPCR primers are described in supplemental Table S7.

### Dual-luciferase reporter assay

The target genes 3’UTR activity was analyzed in both 293T and MDA-MB-231 cells by transient transfection of luciferase reporter constructs. On the 1^st^ day, 6×10^5^/well 293T or 1.5×10^5^/well MDA-MB-231 cells were seeded into 12-well plates. These cells were transfected with 0.17 μ g pmirGLO reporter vector, 1.43 μg pLV-miR-ctrl/pLV-miR-142 vector and 4.0μl lipo-2000 according to manufacturer’s instruction on the next day. 48h after transfection, cell lysates were collected using Passive Lysis Buffer (E1941, Promega). Firefly and Renilla luciferase activity was detected using Dual-Luciferase Reporter Assay System (E1960, Promega) on GloMax^®^-Multi+ Microplate Multimode Reader (Promega).

For gene IL7R and PIK3CG 3kb promoter + 3’UTR reporter analysis, MDA-MB-231 cells were transiently transfected with 0.16 μ g pGL3-IL7R/PIK3CG-3Kb+3’UTR plasmid, 1.36 μ g pLV-miR-ctrl/pLV-miR-142 vector and 0.08 μ g of the control Rluc vector driven by β-actin, TK or CMV promoter, using 4 μl lipo-2000 according to manufacturer’s instruction. Other procedures are the same with 3’UTR dual-luciferase assay.

## Acknowledgements

We greatly appreciate the lab members for valuable discussions. This work was partially funded by NIH grants U01AR069395 (X.Z.) and R01CA172115 and R01CA131231 (P.S.).

## Author contributions

H.T., X.Z. and P.S. conceptualized the study. H.T. performed data analysis and visualization, and wrote the manuscript. S.H. conducted the wet-lab experiments. P.S. and X.Z. revised the manuscript. Z.Z. and X.Q. provided technical assistance. All authors read and approved the final manuscript.

